# MODY mutations in transcription factors *HNF1A* and *HNF1B* affect the production of interferon signaling proteins: evidence at the proteome level

**DOI:** 10.1101/2025.11.24.690114

**Authors:** Ksenia G Kuznetsova, Jakub Vašíček, Dafni Skiadopoulou, Lucas Unger, Rachel Anand Nethala, Michael Wierer, Luiza Ghila, Stefan Johansson, Pål Rasmus Njølstad, Simona Chera, Bente Berg Johansson, Alisa Manning, Marc Vaudel

## Abstract

We investigated the proteomic consequences of MODY-associated mutations in the transcription factors *HNF1A* and *HNF1B* using two cell line models. Quantitative label-free mass spectrometry, employing both data-dependent and data-independent acquisition methods, revealed consistent suppression of energy metabolism and interferon signaling pathways. Pathway and protein interaction analyses confirmed these findings. In cells with the *HNF1A* frameshift mutation, several predicted transcriptional targets, including *A1CF*, were significantly downregulated. These results reinforce the link between HNF1A/HNF1B function and mitochondrial activity as well as innate immune signaling, providing a foundation for further mechanistic studies and causal inference research.

## 1 Introduction

Hepatocyte nuclear factors (HNFs) are a heterogeneous group of DNA-binding transcription factors that are primarily expressed in the liver, but also in other organs such as the pancreas, kidney and intestine. Several HNFs have been implicated in the pathogenesis of diabetes. The protein product of *HNF1A*, hepatocyte nuclear factor 1-alpha (HNF1A), plays a crucial role in glucose homeostasis, beta-cell differentiation, and insulin secretion [1].

Rare pathogenic variants in *HNF1A* represent the most common cause of maturity-onset diabetes of the young (MODY) [2]. In many cases, individuals with HNF1A-MODY can be successfully transitioned from insulin therapy to sulfonylureas — a class of drugs that stimulate insulin secretion — highlighting the essential role of *HNF1A* in this process [3].

*HNF1B* encodes a transcription factor essential for multi-organ development [4]. Pathogenic variants cause HNF1B-MODY, characterized by impaired insulin secretion and pancreatic atrophy, and renal anomalies such as cysts which are a predominant feature. This phenotype is termed as Renal Cysts and Diabetes Syndrome (RCAD) [5]. Common variants in *HNF1A* and *HNF1B* have also been associated with type 2 diabetes (T2D) through genome-wide association studies (GWAS) [6, 7], and rare variants in *HNF1A* through genome and exome sequencing [8]. Despite the complex and heterogeneous nature of the disease, these transcription factors are thus ubiquitous in diabetes pathogenesis.

HNF1A functions as both a homodimer and a heterodimer, binding DNA either as a pair of identical HNF1A molecules or in combination with HNF1B [9]. A frameshift mutation at position 291 in HNF1A, leading to a premature stop codon and production of a truncated protein, is among the most common mutations associated with MODY [10]. This mutation has also been shown to impair the transcriptional regulatory function of both HNF1A and HNF1B by forming non-functional dimers [11].

HNF1A and HNF1B are low-abundance, tissue-specific proteins which makes their study at the proteomic level in human subjects particularly challenging. These proteins are typically not secreted into the bloodstream and can only be detected at low concentrations through tissue biopsy. In this study, we used two cell models to investigate the effects of known MODY-associated mutations on the cellular proteome: *in vitro* generated pancreatic progenitor cells derived from human induced pluripotent stem cells (hiPSCs) carrying the *HNF1A* frameshift mutation (p.291fsinsC, rs587776825), and renal proximal tubule epithelial cells (RPTECs) overexpressing the *HNF1B* S148L point mutation (rs121918674). The effects of these mutations were evaluated at the pathway level to capture broader functional impacts.

## 2 Material & Methods

### 2.1 hiPSC Model of Pancreatic Progenitors with *HNF1A* frameshift mutation

A human induced pluripotent stem cell (hiPSC) model of HNF1A-MODY was created by introducing a frameshift mutation in the transcription factor gene *HNF1A*. The cells were then differentiated up to stage 4, corresponding to pancreatic progenitors. The hiPSCs were purchased from Synthego, the differentiation protocol was adapted from [12], and the p.Pro291fsinsC (p291fsinsC) heterozygous mutation was introduced using the CRISPR-Cas9 method as described in [13]. Control iPSC and HNF1Ap.Pro291fsinsC colonies were maintained in feeder-free maintenance mTeSR Plus cGMP-stabilized medium (StemCell Technologies, 100-0276) and passaged using Gentle Cell Dissociation Reagent (StemCell Technologies, 100-0485) as previously described [14, 15]. All hiPSC cultures were confirmed to be mycoplasma-free using the MycoAlert Mycoplasma Detection Kit (Lonza, LT07-418).

One million cells per sample were harvested, washed with PBS, centrifuged, and frozen at –80°C before being sent for analysis. A total of 12 hiPSC samples were prepared (Fig. 1, Supp. file S2): 3 biological replicates of wild-type cells at stage 0 and stage 4, and 3 biological replicates of mutant cells at both stages. Each replicate underwent independent differentiation. All samples were analyzed by mass spectrometry-based proteomics using both data-dependent acquisition (DDA) and data-independent acquisition (DIA) (see MS method description), resulting in 24 MS data files (Supp. file S2).

**Figure 1.**
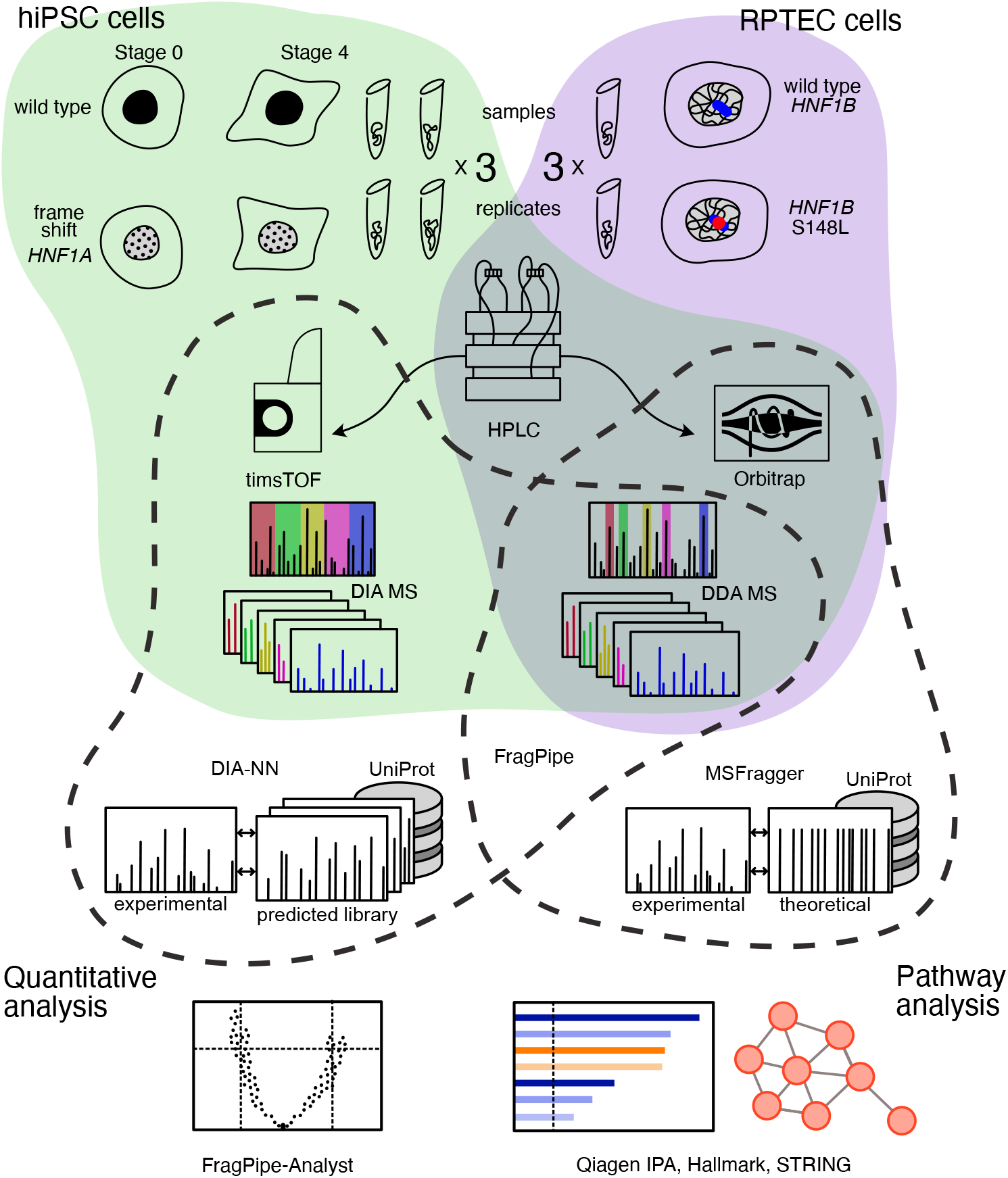
Analysis workflow. A human induced pluripotent stem cell (hiPSC) model of HNF1A-MODY was developed by introducing a frameshift mutation in the *HNF1A* gene. The hiPSCs were differentiated up to stage 4, corresponding to pancreatic progenitors. A total of 12 hiPSC samples were prepared: 3 biological replicates of wild-type cells at stage 0 and stage 4, and 3 biological replicates of mutant cells at both stages. The samples were analyzed using data-dependent acquisition (DDA) and data-independent acquisition (DIA) mass spectrometry modes, resulting in 24 MS data files. An additional experiment was conducted to model HNF1B-MODY using renal proximal tubule epithelial cells (RPTEC/hTERT)). RPTECs were transfected with a plasmid containing the *HNF1B* gene with a point mutation in three biological replicates, resulting in 6 samples, including controls. The samples were processed using high-performance liquid chromatography (HPLC) and analyzed with Orbitrap and timsTOF mass spectrometers. The bioinformatics workflow includes proteomics search, quantitative analysis, and pathway analysis. Data analysis was performed using FragPipe for proteomics searches, and FragPipe-Analyst for quantitative data analysis. Qiagen IPA software, STRING database, and Hallmark gene sets were used for pathway analysis.

### 2.2 RPTECs models with *HNF1B* mutation

Six renal proximal tubule epithelial cell (RPTEC) samples were generated to model HNF1B-MODY. RPTEC/TERT1 (CRL4031™) were obtained from ATCC (United Kingdom). For each sample, 250,000 cells were transfected using Lipofectamine Stem Transfection Reagent (Invitrogen) with the pcDNA3.1-HisB plasmid containing either a point-mutated *HNF1B* construct (S148L) or wild-type *HNF1B*. Cells transfected with the wild-type *HNF1B* vector served as the control. After 24 hours, cells were washed with PBS, harvested, and stored at –80 °C prior to downstream analysis. All conditions were prepared in three biological replicates (Fig. 1, Supp. file S2).

### 2.3 Proteomics sample preparation

Cell pellets were lysed in 50 µL EasyPep lysis buffer (Thermo Fisher Scientific) supplemented with 1 µL Pierce Universal Nuclease (Thermo Fisher Scientific) by pipetting. Samples were incubated for 10 min at 95 °C, and protein concentration was determined using a BCA assay (Pierce). Extracts were reduced with 5 mM TCEP (final concentration) for 15 min at 55 °C, alkylated with 20 mM chloroacetamide (CAA; final concentration) for 30 min at room temperature, and digested with Trypsin/Lys-C at a 1:50 (enzyme:protein) ratio for 16 h at 37 °C. Peptides were desalted using EasyPep peptide clean-up plates (Thermo Fisher Scientific) according to the manufacturer’s instructions. Eluates were dried by vacuum centrifugation, and resuspended in buffer A^*^ (2% acetonitrile, 0.1% TFA).

### 2.4 Liquid chromatography & Mass spectrometry

For data-dependent acquisition analysis (DDA) two hundred nanograms of peptides were loaded onto a 110 cm µPAC NEO HPLC column (Thermo Fisher Scientific) in buffer A (0.1% formic acid in water) using a Vanquish Neo UHPLC system (Thermo Fisher Scientific). Peptides were separated using a four-segment gradient from 5% to 50% buffer B (0.1% formic acid, 80% acetonitrile) in buffer A as follows: 1–5% B over 1 min, 5–10% B over 11.5 min, 10–30% B over 131.5 min, and 30–50% B over 16 min, at a flow rate of 300 nL/min. The separation was followed by column washing and re-equilibration, resulting in a total gradient time of 180 min.

Eluting peptides were introduced via an EASY-Spray source equipped with a 10 µm fused silica emitter (Evosep) into an Orbitrap Ascend Tribrid mass spectrometer (Thermo Fisher Scientific). Data acquisition alternated between a full MS scan (120,000 resolution; scan range, 375–1500 m/z; AGC target, 100%; maximum injection time, Auto) and 20 data-dependent MS/MS scans (15,000 resolution; AGC target, 400%; maximum injection time, 27 ms) with higher-energy collisional dissociation (HCD) activation. The isolation window was set to 1.4 Th, and the normalized collision energy to 25. Dynamic exclusion was applied for 60 s to minimize repeated sequencing of the same precursor ions.

For data-independent acquisition analysis (DIA) peptides were separated on a 25 cm column Aurora Gen2, 1.7 uM C18 stationary phase (IonOpticks) with an EASY-nLC 1200 HPLC (Thermo Scientific) coupled via a captive-spray source to a timsTOF pro2 (Bruker Daltonics) mass spectrometer operated in DIA-PASEF mode [16]. Peptides were loaded in buffer A (0.1% formic acid) and separated with a non-linear gradient of 2 – 35% buffer B (0.1% formic acid, 99.9% acetonitrile) at a flow rate of 400 nL/min over 90 min. Total run time was 101 min including washing phase. The column temperature was kept at 50 °C during all LC gradients. MS acquisition involved 16 diaPASEF scans with two 25 Da windows per ramp, mass range from 400 to 1201 Da, and mobility range of 1.43 to 0.6 1/K0. The collision energy was decreased linearly from 59 eV at 1/K0 = 1.3 to 20 eV at 1/K0 = 0.85. Both accumulation time and PASEF ramp time were set to 100 ms.

### 2.5 Data analysis

Mass spectra were searched against a reference human proteome database downloaded from UniProt in September 2024 consisting of 20,468 entries after the addition of common contaminant sequences. The search was performed by FragPipe 22.0 using a preset “LFQ MBR” workflow for DDA data and “DIA SpecLib DIA PASEF” workflow for DIA data. All DDA and DIA MS files and search engine results are available at ProteomeXchange under the accession numbers PXD069887 and PXD070681 respectively.

The FragPipe output was further statistically analyzed with the FragPipe-Analyst tool [17]. For DDA data the “LFQ” profile was used and the “maxLFQ intensity” was taken as a measure of protein group intensity. For DIA data “DIA” profile was used. We used variance stabilizing normalization method, the “Perseus-type” imputation, and the Benjamini-Hochberg FDR correction implemented in FragPipe-Analyst. The Hallmark gene sets enrichment was run on downregulated proteins. Pathway enrichment and prediction analysis was done with the Qiagen Ingenuity Pathway Analysis (IPA) software (v.134816949). A standard core analysis was run using the data from FragPipe-Analyst filtered to 0.05 Benjamini-Hochberg FDR and log2 fold change absolute values no less than 1.

In order to obtain proteome-level pathway analysis we ran STRING [18] analyses of the differentially abundant proteins identified by FragPipe-Analyst. The proteins were loaded as a list to the multiple proteins analysis, the high confidence (0.7) parameter was chosen for protein interaction identification, and the MCL clustering was implemented with 3 as inflation parameter.

## 3 Results

### 3.1 Depth of analyzed proteomes

We conducted shotgun proteomics on hiPSCs at stage 0 and stage 4 (pancreatic progenitors) using both data-dependent acquisition (DDA) and data-independent acquisition (DIA) methods. At stage 4, we identified: 7,478 proteins using DDA, 7,711 proteins using DIA, with an overlap of 7,013 proteins between the two methods. At stage 0, we identified: 7,737 proteins using DDA, 6,460 proteins using DIA, with an overlap of 6,281 proteins (Supp. file S1-fig.1, Supp. file S3-S7). To trace the effect of the *HNF1A* frameshift mutation on the cellular proteome, we performed quantitative proteomic analysis of differentially abundant proteins by comparing mutant cells to wild-type *HNF1A* controls (Fig. 2). At stage 4, the results were: 315 differentially abundant proteins in the DDA dataset (BH-adjusted p-value *<* 0.05, ≥ 1-fold change), 179 proteins in the DIA dataset, with an overlap of 102 proteins identified in both datasets. At stage 0, the results were: 153 proteins in DDA, 68 proteins in DIA, with an overlap of 9 proteins (Fig. 3, Supp. file S1-fig.1-3, Supp. file S11-S15).

**Figure 2.**
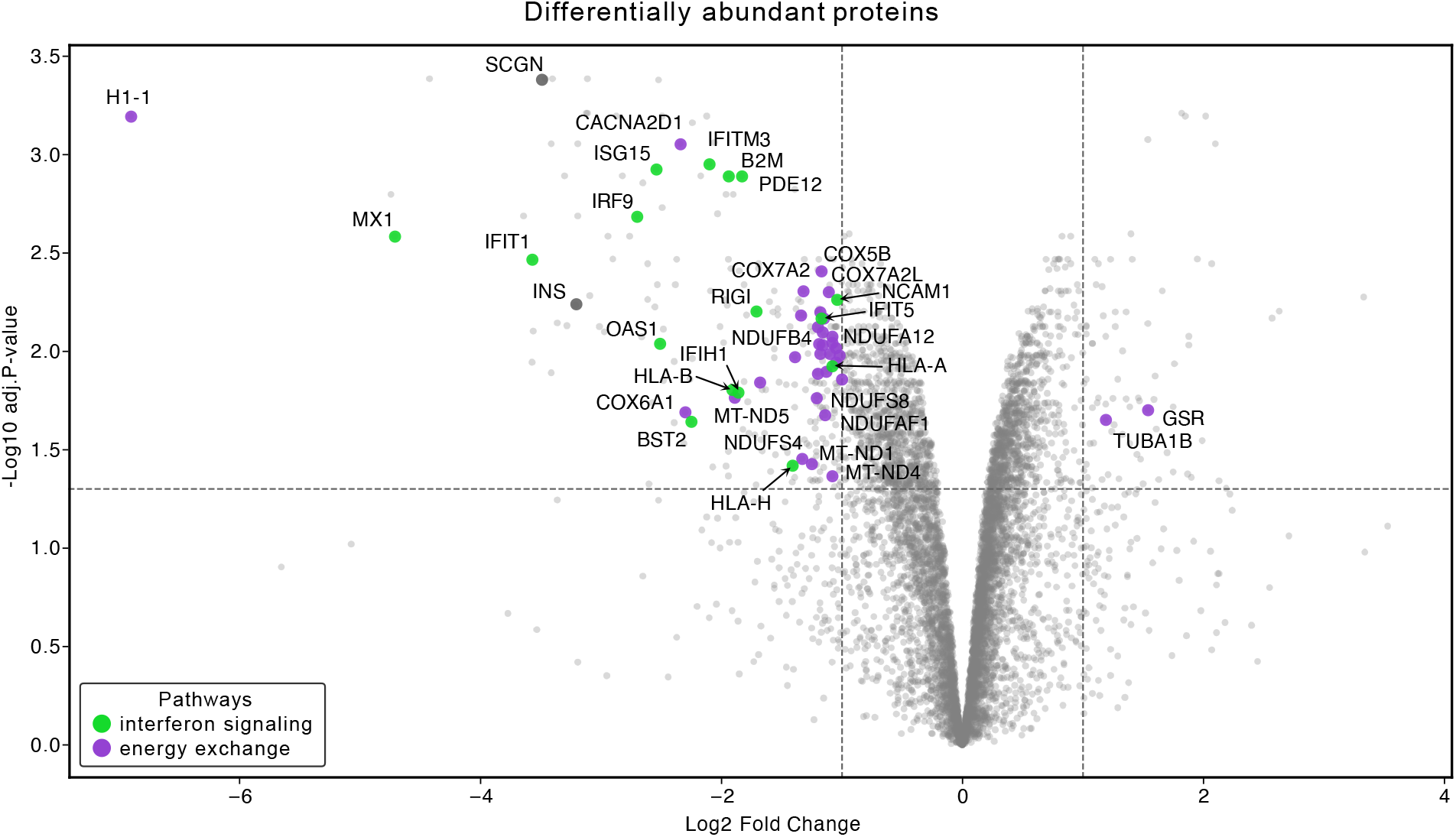
Volcano plot illustrates the differential protein expression in *HNF1A* frameshift mutant hiPSCs compared to wild-type *HNF1A* controls at stage 4, based on data-dependent acquisition (DDA) analysis. The x-axis represents the log2 fold change in protein abundance, while the y-axis represents the -log10 adjusted p-value, indicating the statistical significance of the changes. Proteins with significant upregulation (log2 fold change *>* 1, BH-adjusted p-value *<* 0.05) are shown on the right side of the plot, while proteins with significant downregulation (log2 fold change *<* −1, BH-adjusted p-value *<* 0.05) are shown on the left side.

**Figure 3.**
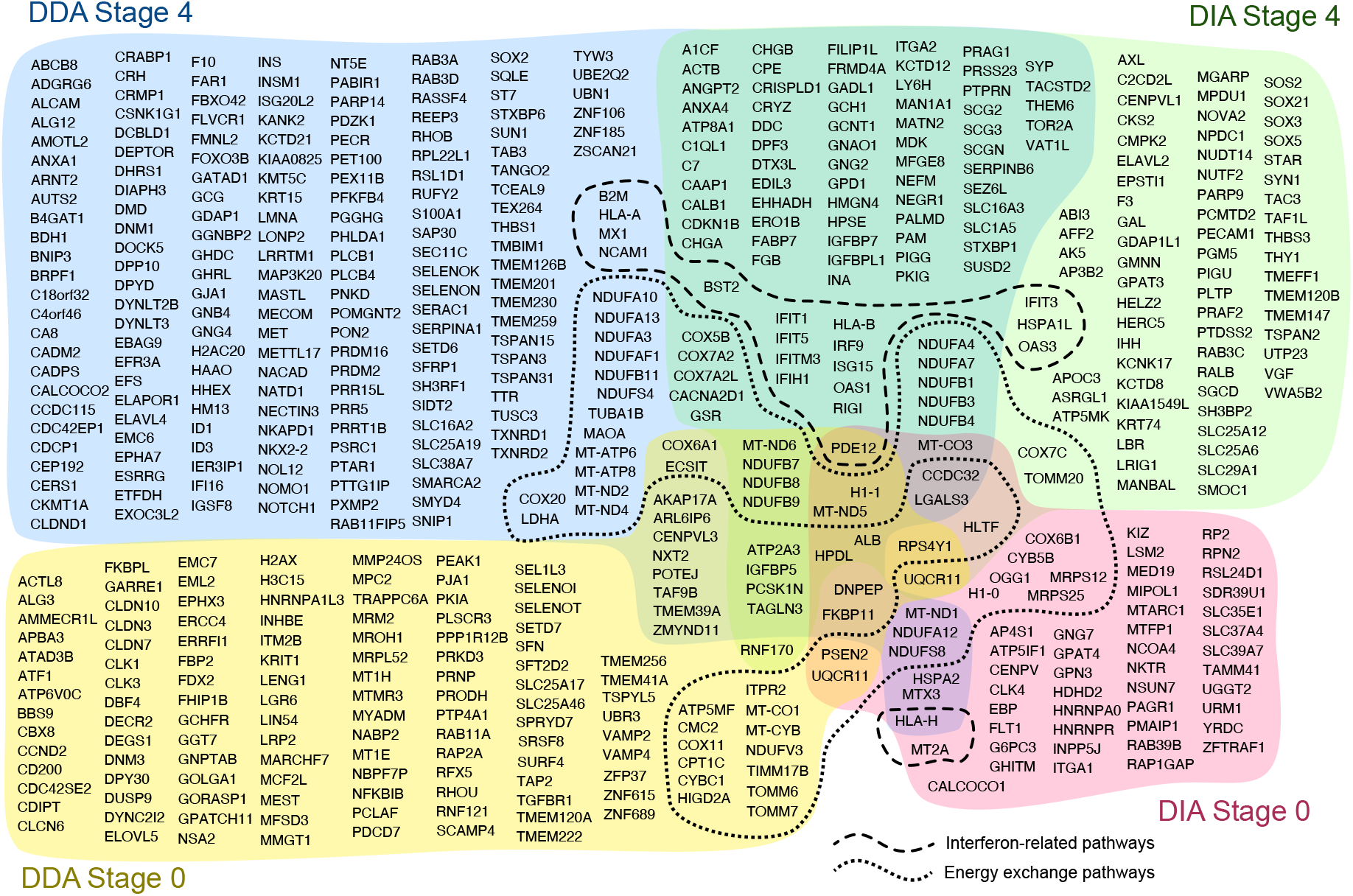
List of differentially abundant proteins in hiPSC. Intersection of results at the individual protein level. Proteins related to energy exchange and interferon signaling pathways (highlighted by dashed lines) as identified by Qiagen IPA.

Analysis of DDA spectra from RPTEC/hTERT samples identified a total of 6,142 proteins (Supp. file S8-S10). Comparison between RPTECs overexpressing *HNF1B* with the S148L point mutation and control cells expressing wild-type *HNF1B* revealed 51 differentially abundant proteins: 30 downregulated and 21 upregulated (Supp. file S1-fig.S9A, Supp. file S16).

### 3.2 Energy and innate immunity pathway disruption

To explore the pathways and biological processes affected by the mutation under investigation, we conducted a core analysis using Qiagen IPA software on all our differentially abundant protein results (Fig. 4, Supp. file S1-fig.S4,5,9B; Supp. file S17-22).

In the hiPSCs at stage 4, the most significantly enriched pathways (B-H adjusted p-value *<* 0.05) that were upregulated (z-score *>* 2) and reproduced in both DDA and DIA analyses of the main dataset were: Granzyme A Signaling, Mitochondrial Dysfunction, Parkinson’s Signaling Pathway, and Sirtuin Signaling Pathway. The significantly downregulated pathways (B-H adjusted p-value *<* 0.05 and z-score *<* −2) included: Oxidative Phosphorylation, Hematoma Resolution Signaling Pathway, Interferon alpha/beta signaling, Interferon Signaling, Interferon gamma signaling, and Cytoprotection by HMOX1. These results suggest that in the cells affected by the frameshift in *HNF1A*, energy metabolism and mitochondrial function are inhibited, along with the innate immune response, particularly interferon signaling. It is important to note that the default core analysis in IPA software evaluates the enrichment of pathways and gene collections from multiple sources, which can lead to an overlap in the reported enriched pathways (Fig. 4A, Supp. file S1-fig.S4A,B; Supp. file S18,20).

**Figure 4.**
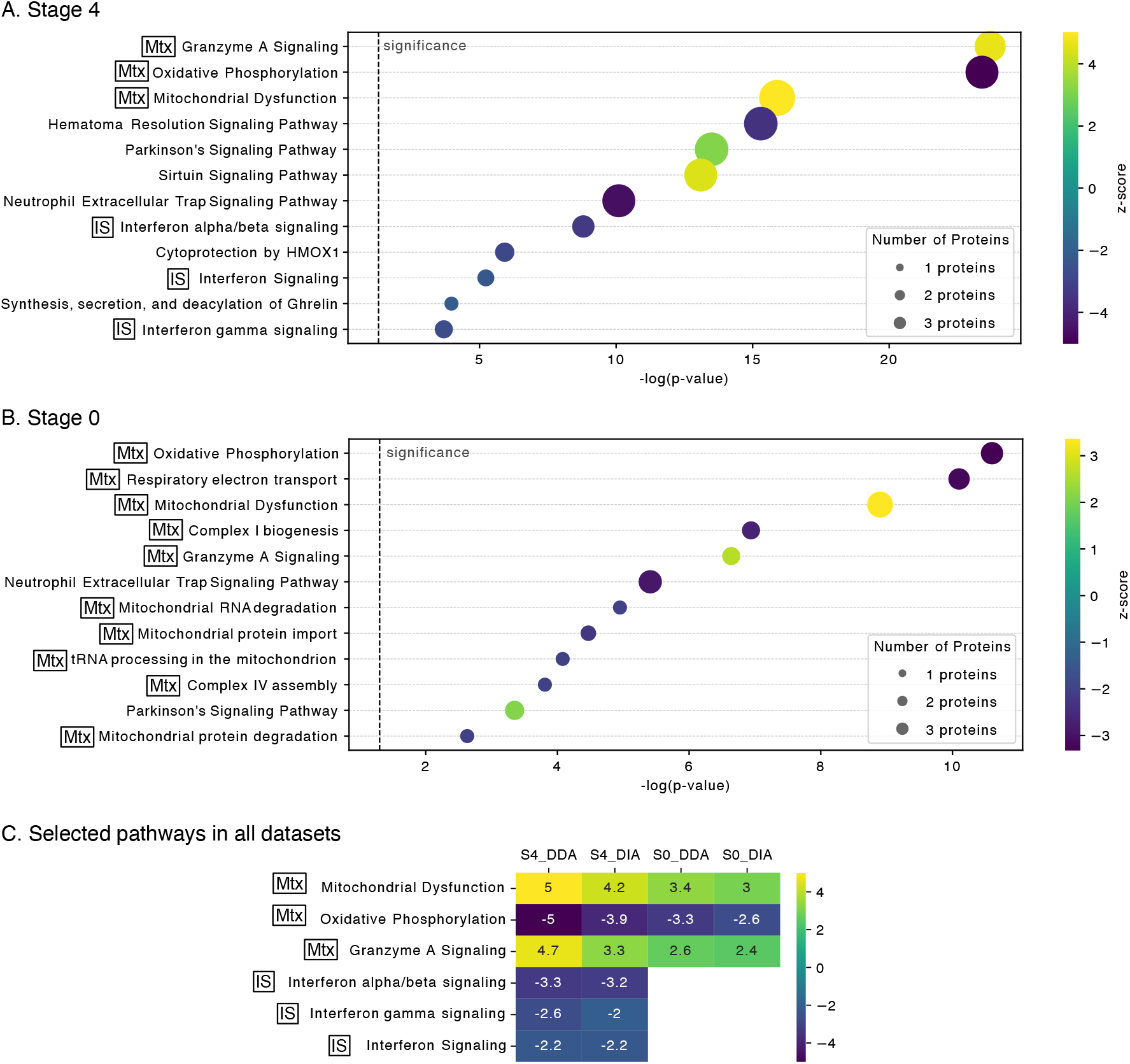
Significantly enriched pathways identified through core analysis using Qiagen IPA software in cells with the *HNF1A* mutation at different stages of differentiation. **A**: Pathways at stage 4, where the most significantly upregulated pathways (B-H adjusted p-value *<* 0.05, z-score *>* 2) include Granzyme A Signaling, Mitochondrial Dysfunction, Parkinson’s Signaling Pathway, and Sirtuin Signaling Pathway. The significantly downregulated pathways (B-H adjusted p-value *<* 0.05, z-score *<* −2) include Oxidative Phosphorylation, Hematoma Resolution Signaling Pathway, Interferon alpha/beta signaling, Interferon Signaling, Interferon gamma signaling, and Cytoprotection by HMOX1. **B**: Pathways at stage 0 in undifferentiated cells. The most significantly upregulated pathways include Mitochondrial Dysfunction, Granzyme A Signaling, and Parkinson’s Signaling Pathway. The downregulated pathways include Oxidative Phosphorylation, Respiratory Electron Transport, Complex I Biogenesis, and Neutrophil Extracellular Trap Signaling Pathway. **C**: Selected pathways across all datasets, highlighting the consistency of the results between DDA and DIA analyses. The pathways are represented by their -log(p-value) and z-score, indicating the level of significance and direction of regulation (up or down). **Mtx**: energy exchange pathways; **IS**: interferon signaling pathways.

At stage 0, in undifferentiated cells, the most significantly upregulated pathways selected by the same criteria were: Mitochondrial Dysfunction, Granzyme A Signaling, and Parkinson’s Signaling Pathway. The downregulated pathways were: Oxidative Phosphorylation, Respiratory Electron Transport, Complex I Biogenesis, and Neutrophil Extracellular Trap Signaling Pathway (Fig. 4B.; Supp. file S1-fig.S4C,D; Supp. file S17,19).

Overall, these results show that the energy exchange pathways were affected in the cells with the mutation already at stage 0, before differentiation towards pancreatic progenitors had occurred. At stage 0 the key pancreatic cell transcription factors such as HNF1A, HNF1B, and HNF4A are not detected by proteomics methods. This is also confirmed in our proteomics data. In contrast, the innate immunity pathways, including interferon signaling, do not show impairment at stage 0 but appear in the results at stage 4 with strong confidence (Fig. 4C, Supp. file S1-fig.S5, Supp. file S21).

In RPTEC pathway analysis using Qiagen IPA showed significant enrichment of Interferon Alpha/Beta Signaling and Interferon Gamma Signaling pathways (Supp. file S1-fig.S9B,C, Supp. file S22). Consistently, analysis of the downregulated proteins against Hallmark gene sets identified enrichment of the Interferon Alpha Response and Interferon Gamma Response pathways. While HNF1A and HNF1B are already known to share functions and interact as transcription factors, our results reinforce this connection and support further investigation into their overlapping regulatory roles. Notably, the specific downregulated proteins in both models strongly overlap, including members of the OAS, IFIT, and HLA families, as well as MX1.

### 3.3 Hallmark enrichment confirms energy exchange and interferon signaling downregulation

As part of the FragPipe-Analyst workflow, we performed enrichment analysis of significantly down-regulated proteins using the Hallmark gene sets [19]. At stage 4, both DDA and DIA analyses revealed strong enrichment in the following gene sets: Oxidative Phosphorylation, Interferon Alpha Response, Interferon Gamma Response, and Pancreatic Beta Cells (Supp. file S1-fig.S6,S9, Supp. file S24-S28). These results suggest potential impairments in energy metabolism, cell differentiation, and innate immune responses. In contrast, at stage 0, the only significantly enriched gene set in both DDA and DIA datasets was Oxidative Phosphorylation. This result confirms that energy exchange pathways are impaired at both differentiation stages, while interferon signaling pathways are only downregulated at stage 4. Thus, the Hallmark gene set enrichment results replicate the findings obtained from Qiagen IPA, reinforcing the conclusion.

### 3.4 Protein interaction networks reflect energy and immune suppression

We analyzed the differentially abundant proteins across all datasets using the STRING database, which is specifically designed for protein-protein interaction analysis. Using the built-in clustering tool, we performed MCL clustering, revealing several significant clusters of differentially expressed proteins (Supp. file S1-fig.S7-8,S10, Supp. file S29-S32). At stage 4, the most enriched clusters with the strongest associations included Oxidative Phosphorylation/Respiratory Electron Transport, consisting of 34 proteins, and Negative Regulation of Viral Genome Replication/Antiviral Defense/Interferon Alpha/Beta Signaling, consisting of 21 proteins. At stage 0, the only enriched cluster observed was Oxidative Phosphorylation/Mitochondrial ATP Synthesis Coupled Electron Transport. These results confirm the effects of the *HNF1A* frameshift mutation at the proteome level, as observed with other pathway analysis tools.

Overall, the regulation of pathways observed at the proteome level using three methods: Hallmark gene sets enrichment, STRING protein interaction analysis and clustering, and core IPA analysis, indicates that the *HNF1A* frameshift mutation is associated with the downregulation of cellular energy exchange processes and the innate immune response.

### 3.5 Proteomics confirms predicted HNF1A and HNF1B targets

From [20] we took the most extensive list of candidate targets and overlapped it with our findings resulting in a list of proteins consisting of 172 everlapped entries. Notably, neither of the most confident targets *COBLL1, NR5A2, HGD, EEF1A1, NHF4A, GPR39* named in the work were found by us in the differentially abundant proteins. On the contrary, in a more recent work investigating the homologous targets of HNF1A in humans and Hnf1a in mice [21], a few genes overlapped with our differentially abundant proteins, including A1CF, proposed as key in HNF1A-mediated glycemia and T2D risk [21]. The other overlapped genes were: *AFF2, ANXA4, DDC, DEPTOR, PGM5, PRR15L, PRRT1B, RAB3D, TTR, VWA5B2*.

The HNF1B predicted target list was taken from JASPAR Predicted Transcription Factor Targets [22]. Out of all candidate targets, *METTL17, NDUFB8* were significantly downregulated in our experiment supporting the hypothesis that HNF1B regulates their expression. The list of overlapped known targets of HNF1A and HNF1B with our findings can be seen in Supp. file S33.

## 4 Discussion

In our study we used label-free mass spectrometry-based proteomics methods to observe the effect of mutations in *HNF1A* and *HNF1B* on the proteome. Proteomics offers limited coverage, yet provides a powerful source of information on protein abundance. Using two acquisition methods, data-dependent (DDA) and data-independent (DIA), we achieved high coverage of proteomes, with over 8,000 identified proteins. The overlap between the datasets produced by the two acquisition methods was, on average, 82% of the number of overall identified proteins. Even though the overlap of the proteins differentially abundant in the comparison between mutant and control cells was lower, the key enriched pathways, Hallmark gene sets, and STRING were consistent. This indicates a mosaic coverage of the proteome capturing similar biological mechanisms (fig. 4, Supp. file S1-fig.S5).

Among the pathways most confidently affected by the frameshift in *HNF1A* were Oxidative Phosphorylation (downregulated), Mitochondrial Dysfunction (upregulated), Granzyme A Signaling (upregulated), and Respiratory Electron Transfer (downregulated), indicating suppression of mitochondrial function and energy exchange. This result aligns with a published proteomics study of human islets, where the authors compared samples with high insulin abundance and low insulin abundance using this parameter as a marker of human islet homeostasis [23]. Mitochondrial dysfunction has also been shown on fully differentiated cells [24]. Insulin itself was also detected and significantly downregulated in our samples with the mutation in *HNF1A* at stage 4 (fig. 2).

Interestingly, energy exchange pathways were already downregulated in both the stem cell stage 0 and stage 4 pancreatic progenitor cells in our analysis, indicating that they might be affected before differentiation occurred and HNF1A protein was produced. Therefore, we observe an association between the *HNF1A* mutation and downregulation of these pathways, although we cannot infer causality between the *HNF1A* and *HNF1B* malfunction and the pathway inhibition. The involvement of *HNF1A* and *HNF1B* in the innate immunity pathways has not been thoroughly investigated, and only sporadic evidence is present in the scientific literature. It has been shown on murine pancreatic islets that, in the context of islet maturation and aging, the expression levels of *Hnf1a* demonstrate the opposite direction of regulation to the interferon signaling pathway [25], and that this pattern is similar between mice and humans. In other words, in developing mice, *Hnf1a* transcription increased while transcription of interferon signaling genes decreased. In aging mice, the decrease in *Hnf1a* transcription was accompanied by an increase in both innate and cellular immune responses. This result may seem to contrast with our findings, where heterozygous mutations presumably leading to partial deactivation of HNF1A and HNF1B led to downregulation of the interferon signaling pathways. This can be explained by the fact that our experiment was conducted on a cell model that did not include the whole heterogeneous islet environment. Indeed, the authors of [25] show that in immunodeficient mice, *Hnf1a* expression remains unchanged upon maturation and aging, indicating that *Hnf1a* modulation is dependent on an immunocompetent environment and not vice versa.

In another study of liver tissue from steatohepatitis patients, the authors also show that the suppression of *HNF1A* is associated with activation of type I interferon signaling [26]. The suggested mechanism in the liver tissue is that HNF1A then acts not as a transcription factor but as a cargo receptor mediating the degradation of TBK1 [26], and acting through another HNF, HNF4A [27], which is a product of a gene also involved in diabetes [28]. We also observe it as downregulated, although it does not reach either the significance or fold change thresholds (Supp. file S12, S14). While these results confirm the observations in mice done in [26], they describe a very different environment of the human liver and cannot be directly compared to our results.

Systemic inflammation has been linked in some studies to *HNF1A* suppression, including a GWAS study showing that genetic variation in *HNF1A* is associated with elevated levels of C-reactive protein in human plasma [29]. Our results confirm the involvement of HNF1A and HNF1B in the immune response, but we observe modulation only in interferon alpha, beta, and gamma signaling, namely the innate immune response.

It is essential to highlight that we are investigating an isolated model of pancreatic beta-cell progenitors and are not observing the effect of the complex islet environment with hormone regulation and various types of cells constituting the islet. Furthermore, in our work, we are investigating the proteomic level, specifically the abundance of proteins rather than the transcription of the genes. While these processes are consequential, they are not always concurrent [30, 31]. Therefore, our results showing suppression of interferon signaling pathways associated with partial deactivation of HNF1A and HNF1B proteins motivate further investigation into their mechanistic relationships. Since many interferon signaling proteins are released into the bloodstream and are measurable with emerging high-throughput proteomics methods, the future direction of research on this question may lie in large patient cohort studies.

We have noticed that linker histone H1-1 was not detected in any of the datasets obtained from the cells with the frameshift in *HNF1A*, compared to the wild-type datasets where the H1-1 signal was high. This effect was reproducible across both differentiation stages and consistent across all datasets analyzed. Another consistently downregulated protein in our stage 4 hiPSC *HNF1A* mutant samples is SCGN (Secretagogin). This protein has been implicated in the pathogenesis of diabetes through its ability to bind insulin directly, contributing to insulin resistance [32]. It has also been proposed as a potential anti-diabetic therapeutic agent [33]. Furthermore, SCGN plays a role in regulating beta-cell proliferation and insulin secretion, processes that are crucial for maintaining glucose homeostasis [34]. Curiously, the *SCGN* and *H1-1* genes are located in close physical proximity (∼365 kb apart) on chromosome 6 but are transcribed from opposite DNA strands. While physical proximity alone does not imply a functional relationship, their simultaneous lower abundance at the proteome level supports the hypothesis of potential co-regulation.

Another notable protein that shows downregulation in the hiPSC stage 4 in both DDA and DIA datasets is APOBEC1 complementation factor (A1CF). This protein has been proposed as an HNF1A target before [21] and shown to regulate alternative splicing of a few transcripts in the beta cells contributing to diabetes pathogenesis. Our proteomic results are in agreement with these findings.

## 5 Conclusions

Using label-free proteomics, we identified that mitochondrial function and interferon signaling pathways are suppressed in hiPSC carrying a p291fsinsC frameshift mutation in the *HNF1A* transcription factor.

Inhibition of interferon signaling became apparent at stage 4 of pancreatic *β*-cell differentiation and is likely a consequence of the malfunctioning HNF1A protein produced due to the frameshift mutation. Although the functional relationships between hepatocyte nuclear factors and interferon signaling proteins remain poorly understood, growing evidence highlights the need for further investigation.

In an additional experiment with RPTEC cells, comparison of cells overexpressing *HNF1B* with the L148S point mutation to control cells overexpressing wild-type *HNF1B* also revealed downregulation of interferon signaling pathways.

Both linker histone H1-1 and secretagogin (SCGN) were less abundant in *HNF1A* frameshift cells; given the physical proximity of their encoding genes, this raises the possibility of their co-regulation.

APOBEC1 complementation factor (A1CF) was downregulated in the cells with a mutation in *HNF1A*, contributing with proteomic evidence to the hypothesis that *A1CF* is the target of this transcription factor and that A1CF is involved in diabetes pathogenesis.

## 6 Data Availability

The mass spectrometry proteomics data have been deposited to the ProteomeXchange Consortium (http://proteomecentral.proteomexchange.org) under the accession numbers PXD069887 for DDA data and PXD070681 for DIA. The code used to analyze the results and create plots is available at https://github.com/kuznetsovaks/fsHNF1Aproteo.git. The supplementary materials are available on Zenodo (https://zenodo.org/records/17672458) under the doi:10.5281/zenodo.17672458.

## 7 Acknowledgments

Mass spectrometry analyses were performed by the Proteomics Research Infrastructure (PRI) at the University of Copenhagen (UCPH), supported by the Novo Nordisk Foundation (NNF) (grant agreement number NNF19SA0059305). The work was supported by the Research Council of Norway (301178), the European Research Council (101171420), and the University of Bergen. The computations were performed on the Norwegian Research and Education Cloud (NREC), using resources provided by the University of Bergen and the University of Oslo (https://www.nrec.no).

## References

[1] M M Byrne, J Sturis, S Menzel, K Yamagata, S S Fajans, M J Dronsfield, S C Bain, A T Hattersley, G Velho, P Froguel, G I Bell, and K S Polonsky. Altered insulin secretory responses to glucose in diabetic and nondiabetic subjects with mutations in the diabetes susceptibility gene MODY3 on chromosome 12. Diabetes, 45(11):1503–1510, November 1996.

[2] K Yamagata, N Oda, P J Kaisaki, S Menzel, H Furuta, M Vaxillaire, L Southam, R D Cox, G M Lathrop, V V Boriraj, X Chen, N J Cox, Y Oda, H Yano, M M Le Beau, S Yamada, H Nishigori, J Takeda, S S Fajans, A T Hattersley, N Iwasaki, T Hansen, O Pedersen, K S Polonsky, and G I Bell. Mutations in the hepatocyte nuclear factor-1alpha gene in maturity-onset diabetes of the young (MODY3). Nature, 384(6608):455–458, December 1996.

[3] Pernille Svalastoga, Alba Kaci, Janne Molnes, Marie H Solheim, Bente B Johansson, Lars Krogvold, Torild Skrivarhaug, Eivind Valen, Stefan Johansson, Anders Molven, Jørn V Sagen, Eirik Søfteland, Lise Bjørkhaug, Erling Tjora, Ingvild Aukrust, and Pål R Njølstad. Characterisation of HNF1A variants in paediatric diabetes in norway using functional and clinical investigations to unmask phenotype and monogenic diabetes. Diabetologia, 66(12):2226–2237, December 2023.

[4] Susan Tucker, Louis Philipson, and Rochelle Naylor. The role of monogenic diabetes in pediatric type 2 diabetes. In Pediatric Type II Diabetes, pages 25–35. Elsevier, 2019.

[5] Coralie Bingham and Andrew T Hattersley. Renal cysts and diabetes syndrome resulting from mutations in hepatocyte nuclear factor-1beta. Nephrol. Dial. Transplant, 19(11):2703–2708, November 2004.

[6] Andrew P Morris, Benjamin F Voight, Tanya M Teslovich, Teresa Ferreira, Ayellet V Segre, Valgerdur Steinthorsdottir, Rona J Strawbridge, Hassan Khan, Harald Grallert, Anubha Mahajan, Inga Prokopenko, Hyun Min Kang, Christian Dina, Tonu Esko, Ross M Fraser, Stavroula Kanoni, Ashish Kumar, Vasiliki Lagou, Claudia Langenberg, Jian’an Luan, Cecilia M Lindgren, Martina Müller-Nurasyid, Sonali Pechlivanis, N William Rayner, Laura J Scott, Steven Wiltshire, Loic Yengo, Leena Kinnunen, Elizabeth J Rossin, Soumya Raychaudhuri, Andrew D Johnson, Antigone S Dimas, Ruth J F Loos, Sailaja Vedantam, Han Chen, Jose C Florez, Caroline Fox, Ching-Ti Liu, Denis Rybin, David J Couper, Wen Hong L Kao, Man Li, Marilyn C Cornelis, Peter Kraft, Qi Sun, Rob M van Dam, Heather M Stringham, Peter S Chines, Krista Fischer, Pierre Fontanillas, Oddgeir L Holmen, Sarah E Hunt, Anne U Jackson, Augustine Kong, Robert Lawrence, Julia Meyer, John R B Perry, Carl G P Platou, Simon Potter, Emil Rehnberg, Neil Robertson, Suthesh Sivapalaratnam, Alena Stančáková, Kathleen Stirrups, Gudmar Thorleifsson, Emmi Tikkanen, Andrew R Wood, Peter Almgren, Mustafa Atalay, Rafn Benediktsson, Lori L Bonnycastle, Noël Burtt, Jason Carey, Guillaume Charpentier, Andrew T Crenshaw, Alex S F Doney, Mozhgan Dorkhan, Sarah Edkins, Valur Emilsson, Elodie Eury, Tom Forsen, Karl Gertow, Bruna Gigante, George B Grant, Christopher J Groves, Candace Guiducci, Christian Herder, Astradur B Hreidarsson, Jennie Hui, Alan James, Anna Jonsson, Wolfgang Rathmann, Norman Klopp, Jasmina Kravic, Kaarel Krjutškov, Cordelia Langford, Karin Leander, Eero Lindholm, Stéphane Lobbens, Satu Männistö, Ghazala Mirza, Thomas W Mühleisen, Bill Musk, Melissa Parkin, Loukianos Rallidis, Jouko Saramies, Bengt Sennblad, Sonia Shah, Gunnar Sigursson, Angela Silveira, Gerald Steinbach, Barbara Thorand, Joseph Trakalo, Fabrizio Veglia, Roman Wennauer, Wendy Winckler, Delilah Zabaneh, Harry Campbell, Cornelia van Duijn, Andre G Uitterlinden, Albert Hofman, Eric Sijbrands, Goncalo R Abecasis, Katharine R Owen, Eleftheria Zeggini, Mieke D Trip, Nita G Forouhi, Ann-Christine Syvänen, Johan G Eriksson, Leena Peltonen, Markus M Nöthen, Beverley Balkau, Colin N A Palmer, Valeriya Lyssenko, Tiinamaija Tuomi, Bo Isomaa, David J Hunter, Lu Qi, Wellcome Trust Case Control Consortium, Meta-Analyses of Glucose and Insulin-related traits Consortium (MAGIC) Investigators, Genetic Investigation of ANthropometric Traits (GIANT) Consortium, Asian Genetic Epidemiology Network-Type 2 Diabetes (AGEN-T2D) Consortium, South Asian Type 2 Diabetes (SAT2D) Consortium, Alan R Shuldiner, Michael Roden, Ines Barroso, Tom Wilsgaard, John Beilby, Kees Hovingh, Jackie F Price, James F Wilson, Rainer Rauramaa, Timo A Lakka, Lars Lind, George Dedoussis, Inger Njølstad, Nancy L Pedersen, Kay-Tee Khaw, Nicholas J Wareham, Sirkka M Keinanen-Kiukaanniemi, Timo E Saaristo, Eeva Korpi-Hyövälti, Juha Saltevo, Markku Laakso, Johanna Kuusisto, Andres Metspalu, Francis S Collins, Karen L Mohlke, Richard N Bergman, Jaakko Tuomilehto, Bernhard O Boehm, Christian Gieger, Kristian Hveem, Stephane Cauchi, Philippe Froguel, Damiano Baldassarre, Elena Tremoli, Steve E Humphries, Danish Saleheen, John Danesh, Erik Ingelsson, Samuli Ripatti, Veikko Salomaa, Raimund Erbel, Karl-Heinz Jöckel, Susanne Moebus, Annette Peters, Thomas Illig, Ulf de Faire, Anders Hamsten, Andrew D Morris, Peter J Donnelly, Timothy M Frayling, Andrew T Hattersley, Eric Boerwinkle, Olle Melander, Sekar Kathiresan, Peter M Nilsson, Panos Deloukas, Unnur Thorsteinsdottir, Leif C Groop, Kari Stefansson, Frank Hu, James S Pankow, Josée Dupuis, James B Meigs, David Altshuler, Michael Boehnke, Mark I McCarthy, and DIAbetes Genetics Replication and Meta-analysis (DIAGRAM) Consortium. Large-scale association analysis provides insights into the genetic architecture and pathophysiology of type 2 diabetes. Nat. Genet., 44(9):981–990, September 2012.

[7] Jacques Murray Leech, Robin N Beaumont, Ankit M Arni, Vignesh Kartik Chundru, Luke N Sharp, Kevin Colclough, Andrew Hattersley, Michael N Weedon, and Kashyap A Patel. Common genetic variants modify disease risk and clinical presentation in monogenic diabetes. medRxiv, February 2025.

[8] Yajie Zhao, Sam Lockhart, Jimmy Liu, Xihao Li, Adrian Cortes, Xing Hua, Eugene J Gardner, Katherine A Kentistou, Marisa Cañadas-Garre, Laurie Fabian, Karen Ho, Nicholas Timpson, Yancy Lo, Jonathan Davitte, David B Savage, Carolyn Buser-Doepner, Ken K Ong, Haoyu Zhang, Robert Scott, Stephen O’Rahilly, and John R B Perry. Population-scale gene-based analysis of whole-genome sequencing provides insights into metabolic health. Nat. Genet., 57(10):2436–2444, October 2025.

[9] D B Mendel, L P Hansen, M K Graves, P B Conley, and G R Crabtree. HNF-1 alpha and HNF-1 beta (vHNF-1) share dimerization and homeo domains, but not activation domains, and form heterodimers in vitro. Genes Dev., 5(6):1042–1056, June 1991.

[10] K Yamagata, Q Yang, K Yamamoto, H Iwahashi, J Miyagawa, K Okita, I Yoshiuchi, J Miyazaki, T Noguchi, H Nakajima, M Namba, T Hanafusa, and Y Matsuzawa. Mutation P291fsinsC in the transcription factor hepatocyte nuclear factor-1alpha is dominant negative. Diabetes, 47(8):1231–1235, August 1998.

[11] Ana-Maria Cujba, Mario E Alvarez-Fallas, Sergio Pedraza-Arevalo, Anna Laddach, Maggie H Shepherd, Andrew T Hattersley, Fiona M Watt, and Rocio Sancho. An HNF1α truncation associated with maturity-onset diabetes of the young impairs pancreatic progenitor differentiation by antagonizing HNF1β function. Cell Rep., 38(9):110425, March 2022.

[12] Alireza Rezania, Jennifer E Bruin, Payal Arora, Allison Rubin, Irina Batushansky, Ali Asadi, Shannon O’Dwyer, Nina Quiskamp, Majid Mojibian, Tobias Albrecht, Yu Hsuan Carol Yang, James D Johnson, and Timothy J Kieffer. Reversal of diabetes with insulin-producing cells derived in vitro from human pluripotent stem cells. Nat. Biotechnol., 32(11):1121–1133, November 2014.

[13] Le Cong, F Ann Ran, David Cox, Shuailiang Lin, Robert Barretto, Naomi Habib, Patrick D Hsu, Xuebing Wu, Wenyan Jiang, Luciano A Marraffini, and Feng Zhang. Multiplex genome engineering using CRISPR/Cas systems. Science, 339(6121):819–823, February 2013.

[14] Shayla Sharmine, Thomas Aga Legøy, Lucas Unger, Joao A Paulo, Luiza Ghila, and Simona Chera. Caloric restriction substantially improves glucose regulation in mice with hnf1a-deficient beta-cells. Acta Physiol. (Oxf.), 241(12):e70121, December 2025.

[15] Lucas Unger, Ulrik Larsen, Shayla Sharmine, Md Kaykobad Hossain, Thomas Aga Legøy, Marc Vaudel, Luiza Ghila, and Simona Chera. Efficient cytoplasmic cell quantification using a semi-automated FIJI-based tool. Sci. Rep., 15(1):27509, July 2025.

[16] Florian Meier, Andreas-David Brunner, Max Frank, Annie Ha, Isabell Bludau, Eugenia Voytik, Stephanie Kaspar-Schoenefeld, Markus Lubeck, Oliver Raether, Nicolai Bache, Ruedi Aebersold, Ben C Collins, Hannes L Röst, and Matthias Mann. diaPASEF: parallel accumulation-serial fragmentation combined with data-independent acquisition. Nat. Methods, 17(12):1229– 1236, December 2020.

[17] Yi Hsiao, Haijian Zhang, Ginny Xiaohe Li, Yamei Deng, Fengchao Yu, Hossein Valipour Kahrood, Joel R Steele, Ralf B Schittenhelm, and Alexey I Nesvizhskii. Analysis and visualization of quantitative proteomics data using FragPipe-Analyst. J. Proteome Res., 23(10):4303–4315, October 2024.

[18] Damian Szklarczyk, Rebecca Kirsch, Mikaela Koutrouli, Katerina Nastou, Farrokh Mehryary, Radja Hachilif, Annika L Gable, Tao Fang, Nadezhda T Doncheva, Sampo Pyysalo, Peer Bork, Lars J Jensen, and Christian von Mering. The STRING database in 2023: proteinprotein association networks and functional enrichment analyses for any sequenced genome of interest. Nucleic Acids Res., 51(D1):D638–D646, January 2023.

[19] Arthur Liberzon, Chet Birger, Helga Thorvaldsdóttir, Mahmoud Ghandi, Jill P Mesirov, and Pablo Tamayo. The molecular signatures database (MSigDB) hallmark gene set collection. Cell Syst., 1(6):417–425, December 2015.

[20] Natasha Hui Jin Ng, Soumita Ghosh, Chek Mei Bok, Carmen Ching, Blaise Su Jun Low, Juin Ting Chen, Euodia Lim, María Clara Miserendino, Yaw Sing Tan, Shawn Hoon, and Adrian Kee Keong Teo. HNF4A and HNF1A exhibit tissue specific target gene regulation in pancreatic beta cells and hepatocytes. Nat. Commun., 15(1):4288, June 2024.

[21] Edgar Bernardo, Matías Gonzalo De Vas, Diego Balboa, Mirabai Cuenca-Ardura, Sílvia Bonas-Guarch, Merce Planas-Felix, Fanny Mollandin, Miquel Torrens-Dinares, Miguel Angel Maestro, Javier García-Hurtado, Sonia Moratinos, Philippe Ravassard, Haiqiang Dou, Holger Heyn, Alexander van Oudenaarden, Nathalie Groen, Eelco de Koning, Christian Conrad, Roland Eils, Santiago Vernia, Patrik Rorsman, and Jorge Ferrer. HNF1A and A1CF coordinate a beta cell transcription-splicing axis that is disrupted in type 2 diabetes. Cell Metab., 37(9):1870–1889.e10, September 2025.

[22] Ieva Rauluseviciute, Rafael Riudavets-Puig, Romain Blanc-Mathieu, Jaime A Castro-Mondragon, Katalin Ferenc, Vipin Kumar, Roza Berhanu Lemma, Jérémy Lucas, Jeanne Cheneby, Damir Baranasic, Aziz Khan, Oriol Fornes, Sveinung Gundersen, Morten Johansen, Eivind Hovig, Boris Lenhard, Albin Sandelin, Wyeth W Wasserman, François Parcy, and Anthony Mathelier. JASPAR 2024: 20th anniversary of the open-access database of transcription factor binding profiles. Nucleic Acids Res., 52(D1):D174–D182, January 2024.

[23] Andreas F Mathisen, Shadab Abadpour, Thomas Aga Legøy, Joao A Paulo, Luiza Ghila, Hanne Scholz, and Simona Chera. Global proteomics reveals insulin abundance as a marker of human islet homeostasis alterations. Acta Physiol. (Oxf.), 239(2):e14037, October 2023.

[24] Fabian L Cardenas-Diaz, Catherine Osorio-Quintero, Maria A Diaz-Miranda, Siddharth Kishore, Karla Leavens, Chintan Jobaliya, Diana Stanescu, Xilma Ortiz-Gonzalez, Christine Yoon, Christopher S Chen, Rachana Haliyur, Marcela Brissova, Alvin C Powers, Deborah L French, and Paul Gadue. Modeling monogenic diabetes using human ESCs reveals developmental and metabolic deficiencies caused by mutations in HNF1A. Cell Stem Cell, 25(2):273–289.e5, August 2019.

[25] Andreas Frøslev Mathisen, Thomas Aga Legøy, Ulrik Larsen, Lucas Unger, Shadab Abadpour, Joao A Paulo, Hanne Scholz, Luiza Ghila, and Simona Chera. The age-dependent regulation of pancreatic islet landscape is fueled by a HNF1a-immune signaling loop. Mech. Ageing Dev., 220(111951):111951, August 2024.

[26] Jinyong He, Cong Du, Xuyun Peng, Weilong Hong, Dongbo Qiu, Xiusheng Qiu, Xingding Zhang, Yunfei Qin, and Qi Zhang. Hepatocyte nuclear factor 1A suppresses innate immune response by inducing degradation of TBK1 to inhibit steatohepatitis. Genes Dis., 10(4):1596– 1612, July 2023.

[27] Cristina Baciu, Elisa Pasini, Marc Angeli, Katherine Schwenger, Jenifar Afrin, Atul Humar, Sandra Fischer, Keyur Patel, Johane Allard, and Mamatha Bhat. Systematic integrative analysis of gene expression identifies HNF4A as the central gene in pathogenesis of non-alcoholic steatohepatitis. PLoS One, 12(12):e0189223, December 2017.

[28] E R Pearson, S Pruhova, C J Tack, A Johansen, H A J Castleden, P J Lumb, A S Wierzbicki, P M Clark, J Lebl, O Pedersen, S Ellard, T Hansen, and A T Hattersley. Molecular genetics and phenotypic characteristics of MODY caused by hepatocyte nuclear factor 4alpha mutations in a large european collection. Diabetologia, 48(5):878–885, May 2005.

[29] Angela D Armendariz and Ronald M Krauss. Hepatic nuclear factor 1-α: inflammation, genetics, and atherosclerosis. Curr. Opin. Lipidol., 20(2):106–111, April 2009.

[30] Komal P Singh, Christine Miaskowski, Anand A Dhruva, Elena Flowers, and Kord M Kober. Mechanisms and measurement of changes in gene expression. Biol. Res. Nurs., 20(4):369–382, July 2018.

[31] Christopher Buccitelli and Matthias Selbach. mRNAs, proteins and the emerging principles of gene expression control. Nat. Rev. Genet., 21(10):630–644, October 2020.

[32] Anand Kumar Sharma, Radhika Khandelwal, M Jerald Mahesh Kumar, N Sai Ram, Amrutha H Chidananda, T Avinash Raj, and Yogendra Sharma. Secretagogin regulates insulin signaling by direct insulin binding. iScience, 21:736–753, November 2019.

[33] Anand Kumar Sharma, Radhika Khandelwal, Swathi Chadalawada, N Sai Ram, T Avinash Raj, M Jerald Mahesh Kumar, and Yogendra Sharma. SCGN administration prevents insulin resistance and diabetic complications in high-fat diet fed animals. bioRxiv, September 2017.

[34] Shuhui Ouyang, Sunmin Xiang, Xin Wang, Xin Yang, Xuan Liu, Meilin Zhang, Yiting Zhou, Yang Xiao, Lingzhi Zhou, Gang Fan, and Jing Yang. The downregulation of SCGN induced by lipotoxicity promotes NLRP3-mediated β-cell pyroptosis. Cell Death Discov., 10(1):340, July 2024.

